# State aggregation for fast likelihood computations in molecular evolution

**DOI:** 10.1101/035063

**Authors:** Iakov I. Davydov, Marc Robinson-Rechavi, Nicolas Salamin

**Affiliations:** Department of Ecology and Evolution, Biophore, University of Lausanne, 1015 Lausanne, Switzerland; Swiss Institute of Bioinformatics, Genopode, Quartier Sorge, 1015 Lausanne, Switzerland

## Abstract

**Motivation:** Codon models are widely used to identify the signature of selection at the molecular level and to test for changes in selective pressure during the evolution of genes encoding proteins. The large size of the state space of the Markov processes used to model codon evolution makes it difficult to use these models with large biological datasets. We propose here to use state aggregation to reduce the state space of codon models and, thus, improve the computational performance of likelihood estimation on these models.

**Results:** We show that this heuristic speeds up the computations of the M0 and branch-site models up to 6.8 times. We also show through simulations that state aggregation does not introduce a detectable bias. We analysed a real dataset and show that aggregation provides highly correlated predictions compared to the full likelihood computations. Finally, state aggregation is a very general approach and can be applied to any continuous-time Markov process-based model with large state space, such as amino acid and coevolution models. We therefore discuss different ways to apply state aggregation to Markov models used in phylogenetics.

**Availability:** The heuristic is implemented in the `godon` package (https://bitbucket.org/Davydov/godon) and in a version of FastCodeML (https://gitlab.isb-sib.ch/phylo/fastcodeml).

## Introduction

Evolutionary models are necessary to study the processes governing the evolution of genes, genomes and organisms. While relatively simple models are often sufficient to provide a good estimation of species or gene trees, inferring the specific processes that govern the evolution of molecular data (e.g. selection or co-evolution) requires more complex models. The ability to apply these complex models to large datasets involving many genes and/or species offers the promise to better understand evolution in a more general context. This approach has, however, an important computational cost because of the large numbers of parameters and/or the large size of the state space involved in these complex models.

The computational performance of phylogenetic methods has always been an important issue in molecular evolution. Likelihood-based methods in phylogenetics would not be possible without the use of Felsenstein’s tree pruning algorithm (Felsenstein, 1981) coupled with the growth of computer performance. However, these methods only became commonly used with the heuristics implemented in software such as PhyML and RaXML (Guindon *et al.,* 2010; Stamatakis, 2014). Recent years have thus seen tremendous decreases in computing times, to the extent that data sets with thousands of sequences can now be analysed. However, most progress has been made on simple models of DNA or amino-acid evolution. More complex models, such as codon models used to detect selection, are still computationally too costly to be applied on large genomic datasets (e.g. all Ensembl Compara; Vilella *et al.* 2009).

The complexity of codon models comes from the large state-space that is necessary to represent the 61 codons (excluding the three stop codons). The simplest codon model, which is called M0 (Goldman and Yang, 1994), assumes a single parameter *ω* to model a constant selective pressure occurring on all sites and branches of a phylogenetic tree. The M0 model is probably not realistic enough and more complex models that involve multiple transition matrices have been developed to detect episodic positive selection on a subset of sites and of phylogenetic branches (Zhang *et al.,* 2005; Murrell *et al.,* 2012; Smith *et al.,* 2015). One of the most commonly used complex models is the branch-site model (Zhang *et al.,* 2005), which assumes three classes of selection (parameters *ω*_0_, *ω*_1_, *ω*_2_ with *ω*_2_ allowing positive selection) on sites along specific branches of the tree (called foreground branches) and two classes (parameters *ω*_0_ and *ω*_1_) on the other branches.

Since an accurate phylogenetic tree is critical to evolutionary and comparative studies, most developments to speedup the parameter estimation of evolutionary models have focused first on the optimization of search strategies to find the tree topology and branch lengths. Examples include the choice of the starting tree topology (Huelsenbeck *et al.,* 2001; Guindon and Gascuel, 2003; Stamatakis *et al.,* 2004; Stamatakis, 2014; Nguyen *et al.,* 2014), improved tree rearrangements strategies (Swofford and Olsen, 1990; Guindon and Gascuel, 2003; Hordijk and Gascuel, 2005; Stamatakis *et al.,* 2005; Nguyen *et al.,* 2014), computation economy (Goloboff, 1993; Gladstein, 1997; Ronquist, 1998), and independent branch-length estimation (Guindon and Gascuel, 2003).

However, an important part of the computational cost is spent calculating the likelihood function itself. Although this part is not the most limiting step for tree searching methods using simple models, it becomes a major bottleneck for the evaluation of more complex evolutionary scenarios such as codon models. In this case, the reuse of the eigenvectors and eigenvalues for a set of branches can improve computational performance (Schabauer *et al.,* 2012; Valle *et al.,* 2014). Other optimization techniques that involve, for example, transforming the problem of exponentiating an asymmetric matrix into a symmetric problem, or performing matrix-matrix multiplication rather than matrix-vectors for the estimation of conditional vectors, have also been shown to speedup the calculations of the likelihood (Schabauer *et al.,* 2012). There has also been some progress on Bayesian computation, e.g. using data augmentation (Lartillot, 2006; Rodrigue *et al.,* 2008; de Koning *et al.,* 2012). Despite these improvements, likelihood calculations still remain computationally intensive.

The size of the state-space of the continuous-time Markov chain directly impacts the most computationally intensive steps of this likelihood computation, since it affects the size of the rate and probability matrices (*Q* and *P*, see below), as well as of the conditional probability vector. A method allowing a reduction of the number of states while affecting minimally the precision of the likelihood estimation is therefore a potentially interesting avenue to further reduce the computational burden of these methods.

We propose here a heuristic method to speedup matrix exponentiation and partial likelihood calculations by reducing the number of states in a continuous-time Markov chain without losing the complexity of the model. We use state aggregation techniques to selectively combine states of the instantaneous rate matrix. We illustrate this technique with a simple and a complex codon model, since their state-space is relatively large (61 states). We show using simulations and the analysis of an empirical dataset that aggregation can provide significant speedup for codon models, with a very low cost in terms of accuracy. We further discuss the potential biological applications that could benefit from this approach to illustrate the wide applicability of state aggregation.

### Key steps of likelihood computation in phylogenetics

The performance of the likelihood calculations are governed by two computationally intensive steps: matrix exponentiation and matrix-vector multiplication.

Matrix exponentiation is at the heart of models based on continuous-time Markov chains. The rate of change from one state to any other in an infinitesimally small time interval is given by the instantaneous rate matrix *Q*. The probability of changing between the states of the process in a time interval *t* is then given by the probability matrix *P*: *P*(*t*) = *e*^*Qt*^. For computational purposes, the rate matrix is first diagonalized such that *Q* = *U*Λ*U*^−1^, where *U* is the matrix of eigenvectors and Λ is a matrix whose diagonal elements correspond to the eigenvalues of the instantaneous matrix *Q*. This matrix decomposition allows the probability matrix *P* to be quickly computed for any time interval *t* as *P*(*t*) = *e*^*Qt*^ = *Ue*^*Λt*^*U*^−1^.

Branches of a phylogenetic tree represent the evolutionary path between an ancestral sequence and its descendants. We therefore need to compute the matrix *P* for every branch of a tree. The instantaneous rate matrix *Q* needs thus to be exponentiated for every branch length. The probabilities of observing the states in the ancestral sequence are then calculated by multiplying the conditional probability vectors for each descendant branch. These probability vectors are obtained by multiplying the *P* matrix for branch *i* with the conditional vector of the corresponding descendant. This procedure, known as Felsenstein’s tree pruning algorithm, is repeated for every node of the phylogenetic tree until we reach the root of the tree (Felsenstein, 1973).

## Algorithm

### State Aggregation

The computational cost of the two steps described above highly depends on the state-space of the continuous-time Markov chain used. Any reduction in the state-space can therefore increase the efficiency of the likelihood calculations. We investigate here the use of state aggregation to combine states of a Markov chain into several groups and therefore reduce the complexity of matrix exponentiation and matrix-vector multiplication.

Let us consider a Markov chain taking values in a finite set *S* = {*A*_1_, *A*_2_,…, *A*_n_} with transition matrix *P* and stationary frequencies *π*_1_, *π*_2_,…, *π*_*n*_. Let *S*^*c*^ = {*A*_1_, *A*_2_,…, *A*_*m*_} be a set of states to be aggregated, where *m* < *n*.

The aggregated chain will have a space of

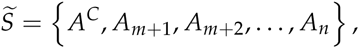

where *A*^*C*^ is the aggregated state. The new aggregated state *A*^*C*^ changes the entries of the probability matrix *P* in the following way:

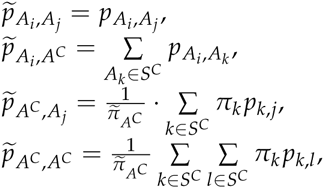

where *A*_*i*_, *A*_*j*_ ∉*S*^*C*^.

The stationary frequencies are estimated as 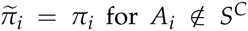. These stationary frequencies are consistent with frequencies of the original Markov chain. The frequencies of the aggregated state is estimated as 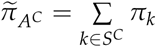.

The same method can be applied at the level of the instantaneous rate matrix *Q*. The diagonal elements of the matrix must however be set to 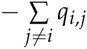 to ensure that the sum of every row is equal to zero (Fig. S1 B, C) (Aldous and Fill, 2002, chapter 2).

### Aggregation for Codon Models

An obvious question in performing aggregation is the definition of “similar states” to aggregate. We define all non observed states for a position of the alignment to be “similar” in the context of that position. The rationale is that the codons that are not observed at this site in any of the sequences at the tips of the tree have low probability to occur as ancestral states. The lack of some possible codons could be due to chance, but in many cases we expect a subset of codons to occur at a site because of natural selection or mutational bias. For example, a protein site which is constrained to be negatively charged will only use codons encoding such amino acids. It is thus justified to call all other codons “similar” relative to this site. We therefore aggregated all states unobserved at a position (i.e. triplet of columns of the DNA alignment) into a single state (Fig. 1). The approach that we use here to aggregate states in codon models resembles the models of amino acid and nucleotide substitutions proposed in Yang *et al.* (1998); Susko and Roger (2007); Phillips *et al.* (2004); Vera-Ruiz *et al.* (2014). However, we propose to select the new aggregated state-space independently for each position of the alignment, which was not done in the amino acid and nucleotide contexts. Note that we performed in this study only the aggregation on the probability matrix *P*. We discuss the advantages of aggregation on the *P* or *Q* matrices in the Discussion section.

**Figure 1:**
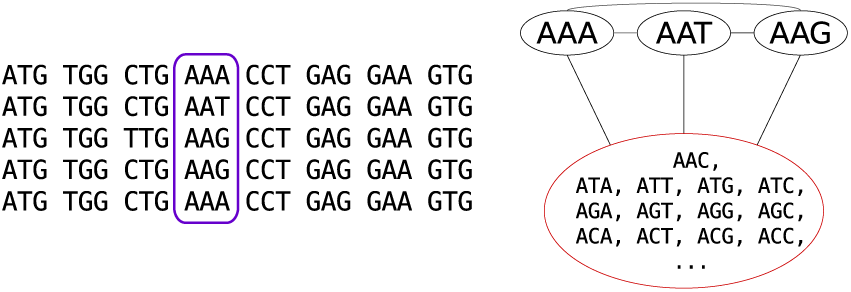
Example of state aggregation for one position (highlighted in purple) in a codon alignment.

For the aggregated process to have a Markovian property, it has to satisfy the lumpability condition, i.e. *pA*_*i*_,*A*_*k*_ = *pA*_*j*_,*A*_*k*_ should be true for any *i*, *j* ∈ *A*^*C*^ and 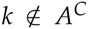 (Kemeny and Snell, 1983), or equivalently for the instantaneous rate matrix (Hillston, 1995). This condition is generally not satisfied with respect to an arbitrary aggregation scheme, as this would require all the transition rates or substitution probabilities to have the same value. Moreover, in the widely used codon substitution models, double substitutions are not allowed and their respective transition rates are set to zero, which makes lumpability condition unsatisfiable. Thus the proposed technique should be viewed as a heuristic.

The intensity of state aggregation can be modified and we tested three different approaches by implementing them for the codon model M0 (Goldman and Yang, 1994). The first and least aggressive approach aggregates only the positions that were absolutely conserved in any sites of the alignment. The state-space for these sites is thus reduced to two states: the observed (conserved) codon, and the “meta-state” of the 60 other non stop codons. In the second approach, all positions were aggregated and the “meta-state” included all codons not present in the position subjected to aggregation. In the third approach, all the positions were aggregated, but the “meta-state” included only codons corresponding to the amino-acids not present at the current position. This can be viewed as a less-aggressive version of the second approach utilizing properties of the genetic code. The first two approaches represent extreme cases of the application of aggregation, while the third one is more moderate.

Given the small speedup of the first and third approaches on M0 (see Results), only the second approach was employed for the more complex branch-site model.

Additionally, two random aggregation strategies were evaluated. These strategies were used as a control to determine if our choice of state partitioning is better than random. In the first strategy, “meta-states” of full aggregation were shuffled between the alignment positions. This should give a speedup similar to the full aggregation, while not relying on codons present at each position. In the second random strategy, the state-space was randomly split into “meta-states”, while keeping the total number of states per position. The number of states stays the same in this case, but the computations are expected to take more time since multiple “meta-states” are present.

## Materials and Methods

### Software

State aggregation for the M0 model was implemented in the `godon` package (https://bitbucket.org/Davydov/godon). We selected an optimization algorithm with a large but fixed number of iterations (10,000 iterations in this case) to reduce the influence of random factors associated with the optimization trajectory on the total computation time.

State aggregation for the branch-site model was implemented in a version of `FastCodeML` (https://gitlab.isb-sib.ch/phylo/fastcodeml, branch `agg`), which is a software that has been optimized for computational efficiency of the calculation of the matrix exponentiation and the matrix-vector multiplication (Valle *et al.,* 2014).

All sequence simulations were performed using the evolver program from the PAML package (Yang, 2007).

### Dataset

Six datasets were simulated for the M0 model (see Table S1). We varied one parameter at a time, based on the following settings: 300 codons, 18 sequences, *ω*_0_ = 0.3, *κ* = 2, equal codon frequencies (*π*_*i*_ = 1/61), default tree length (4). We used the the ENSGT00680000099620 gene tree from the Ensembl database (Cunningham *et al.,* 2015) for topology and relative branch lengths.

For the branch-site model, 2,000 alignments were simulated with stochastic birth-death trees and *κ*, *ω*_0_, *ω*_2_, *p*_0_, *p*_1_, alignment length and number of tips sequences chosen randomly (Table S2 A, Fig. S2). One thousand of the alignments were simulated under the branch-site model null hypothesis with *ω*_2_ = 1, while the other 1000 alignments represented the alternative hypothesis with *ω*_2_ > 1. In these simulations every parameter was drawn randomly from a specific distributions (Table S2) to obtain more biologically realistic datasets. We chose at random a single foreground branch to perform the simulations and the same foreground branch was used foreground the inferrence. We used `evolver` from `PAML` 4.8 for simulation (Yang, 2007).

We also simulated datasets using an *extended branch-site model*. In this model, *ω*_0_ and *ω*_2_ (when *ω*_2_ > 1) were replaced by a set of discrete categories created from *Beta* and *Gamma* distributions respectively (five discrete categories were used). We also incorporated *Gamma* distributed site-rate variations (Rubinstein *et al.,* 2011). The parameters for these distributions are described in Table S2 B and Fig. S3. We used the `cosim` package for the simulations (https://bitbucket.org/Davydov/cosim).

The likelihood ratio test (LRT) was used for model selection, with a significance level of α = 0.05.

We performed multiple hypothesis testing correction using the `qvalue` R package, *π*_0_ was estimated using the bootstrap method (Storey *et al.,* 2004).

Finally, a Primates dataset from the Selectome database (Proux *et al.,* 2009; Moretti *et al.,* 2014) release 6 was used to study the behaviour of the method on a real dataset. The dataset consists of 15669 gene trees and alignments (http://selectome.unil.ch/cgi-bin/download.cgi). We tested the inferrence of selection on every non-terminal branch of the Primates trees.

## Results

### M0 Model

For the simple M0 model, we first compare the performance of likelihood maximization in three different modes: full likelihood (no aggregation), aggregation for conserved positions and full aggregation. Here we kept the branch lengths fixed and optimized *ω* and *κ*.

The parameter values obtained for all datasets using both aggregation modes are highly correlated with values estimated by the full likelihood (Fig. S4, S5, S6, S7, S8, S9). The error in estimation of the parameters is small and is not dependent on the simulation parameters (Fig. 2, S10, S11), with the exception of tree length (Fig. 3). The bias in parameter estimation associated with the long trees is smaller for less aggressive aggregation strategies (Fig. S12). Comparisons with the two random aggregation strategies show noticeably better accuracies in parameter estimation with the observation-based aggregation (Fig. S13).

**Figure 2:**
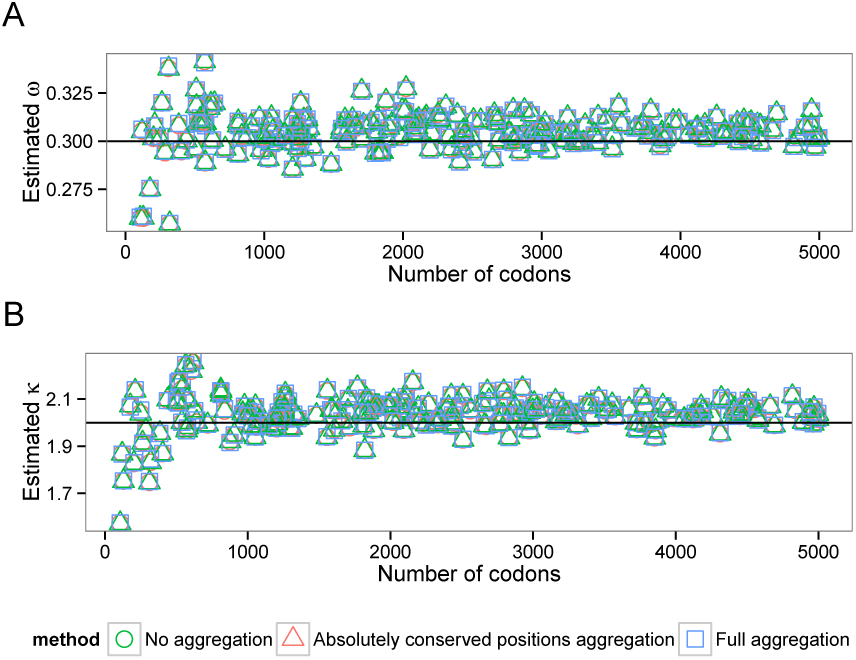
Estimated ω(A) and κ(B) values depending on the alignment length (`alen` dataset, M0 model). Lines correspond to the simulation parameter values.

**Figure 3:**
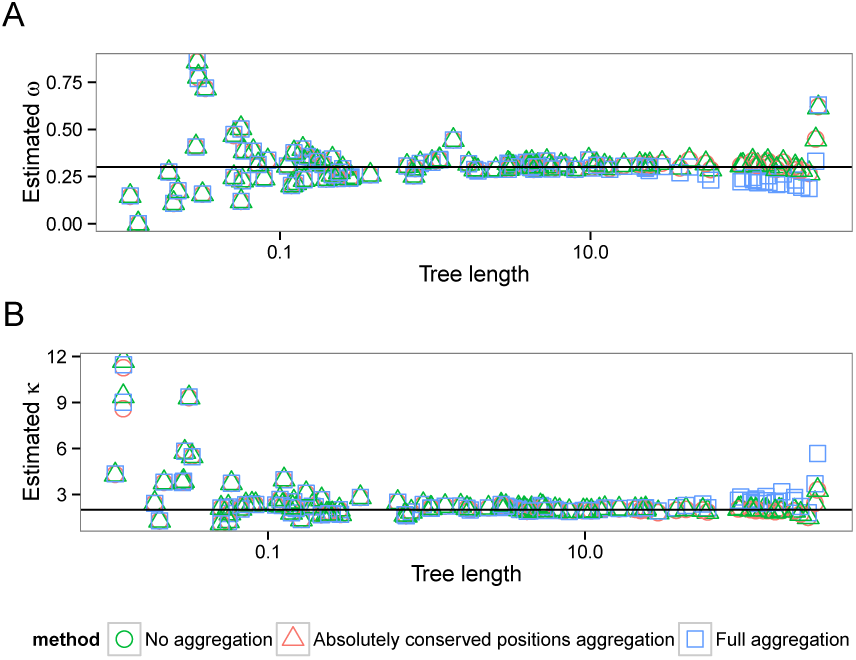
Estimated ω(A) and κ(B) values depending on the tree length (`tlen` dataset, M0 model). Lines correspond to the simulation parameter values. Tree length limited to the range [0.01;300], see text).

The mean computational speedup is approximately 1.7 for aggregation on all positions (Fig. 4), but only 1.2 and 1.02 (Fig. S14) for genetic code based aggregation and aggregation limited to fixed positions respectively. We thus only analyzed in details the behavior of the full aggregation mode.

**Figure 4:**
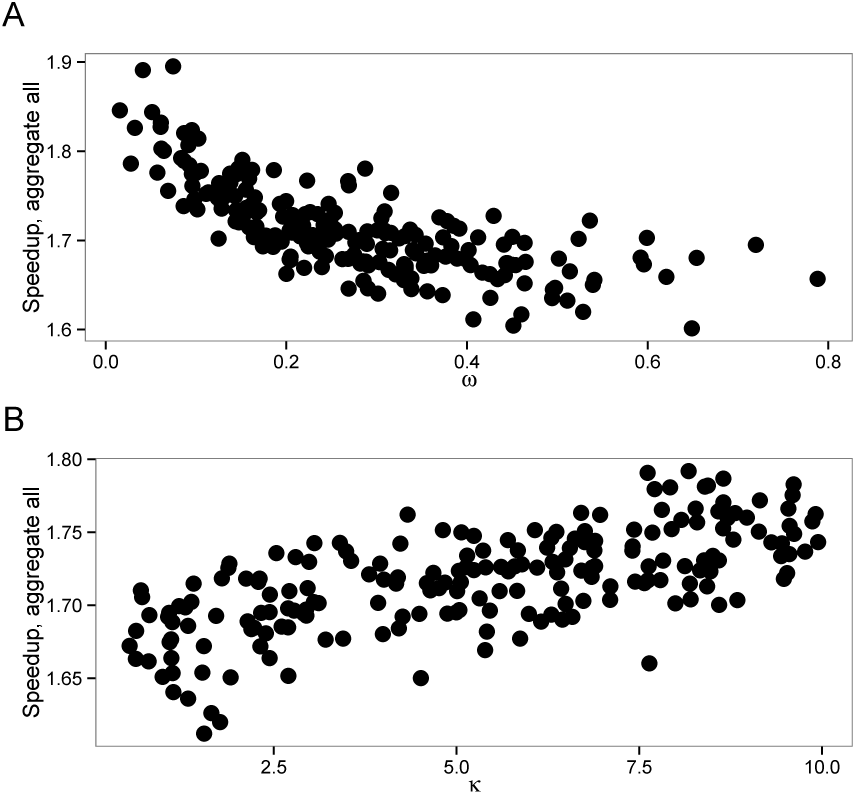
Speedup of aggregation on all alignment positions with M0 model. A) `wvar` dataset with variable ωvalue, B) `kvar` dataset with variable κ.

First, we see a strong effect of the alignment length on the speedups obtained (Fig. 5). Matrix eigendecomposition is performed only once per likelihood evaluation and the decrease in the state-space between the full likelihood and the aggregation does not have any impact on the eigendecomposition performance. However, a longer alignment will increase the number of times the tree pruning step is performed (i.e. once per site), which becomes more important in the overall computational cost. For instance, the speedup obtained with an alignment of 500 codons is 1.8 with full aggregation. The maximum speedup of 6.8 fold was achieved on extremely long alignments (above 10,000 codons) and short trees (total length < 0.05).

**Figure 5:**
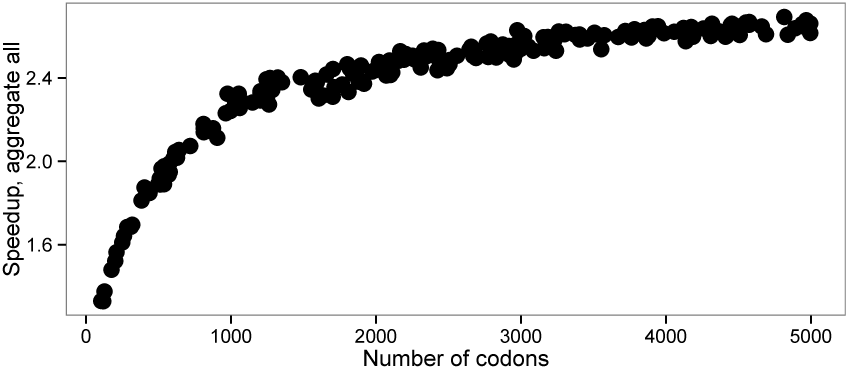
Speedup depending on a number of the codons (`alen` dataset, M0 model).

While there is a larger error on the estimation of model parameters (*κ* and *ω*) with shorter alignments, this effect is identical with or without aggregation (Fig. 2). The heuristic that we propose does therefore not increase error on a simple model even with short alignments. Interplay between eigendecomposition and pruning times explains the direct effect of the relationship between the number of sequences and the speedup (Fig. 6). A large number of sequences decreases the proportion of time spent in the eigendecomposition phase and subsequently increases the speedup.

**Figure 6:**
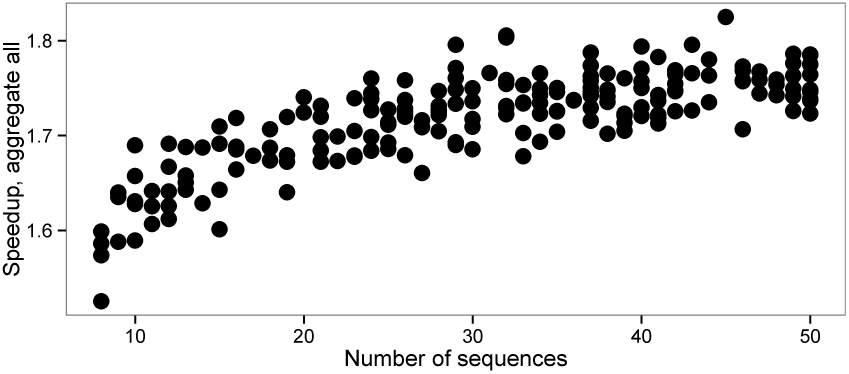
Speedup versus number of sequences (`nseq` dataset, M0 model).

Changes in the other parameters impact the speedup of the aggregation mode insofar as they change the number of codon states per alignment site. The latter has then a direct effect on the number of non aggregated states. Indeed, we aggregate into one state all codons which are not observed in a given position. Thus any processes that reduces the number of different codons per position also increases the efficiency of aggregation. Hence, the speedup is slightly higher for smaller *ω* values because, as *ω* approaches 0, more and more codons at a particular site are only part of a synonymous codon set. The number of possible codons is thus greatly reduced and there is a higher chance that the aggregation will lead to very few states. In contrast, increasing *ω* values will lead to an increasing number of states observed. Similarly, extremely short branches limit state variety at each site, which in turn increase the level of aggregation possible and thus increase speedup (Fig. 7). Biased codon frequencies can also reduce diversity of states and thus increase aggregation speedup. In our simulations, codon frequencies were drawn from a Dirichlet distribution and we varied the concentration parameter *α* to estimate the effect of codon frequencies on codon aggregation. We see a better speedup associated with smaller values of the *α* parameter, which leads to a higher variance between codon frequencies (Fig. S15).

**Figure 7:**
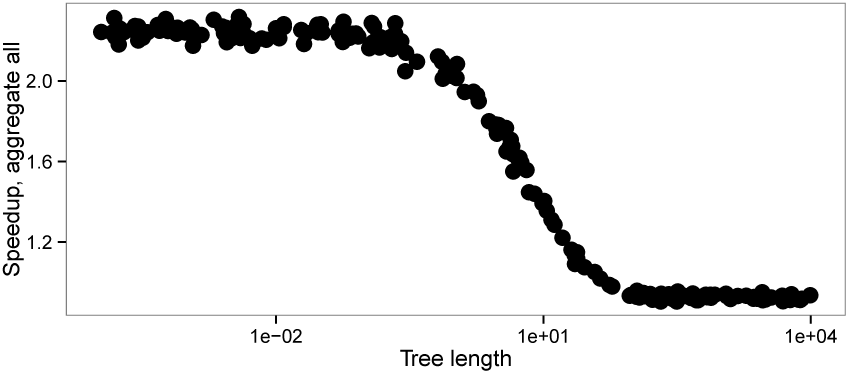
Speedup versus tree length (`tlen` dataset, M0 model).

Total tree length is the only parameter in our simulations that also affects the accuracy of the estimation of model parameters (Fig. 3, S16). Longer trees tend to improve the accuracy of the estimation of the parameters *ω* and *κ*. However, extremely long trees lead to an increase in error both in aggregated and in full likelihood mode, probably because of saturation. It appears that under reasonable conditions of applicability of the M0 model (i.e. total tree length <20 substitutions per codon), aggregation does not lead to any detectable bias, while for extremely long trees aggregation can introduce a slight bias.

We also estimated the branch lengths during the optimization of the M0 model. There was no systematic bias in branch lengths estimation for short trees (Fig. S17), while we observed an increased error in branch lengths estimation on extremely long trees (Fig. S18). The error on branch lengths was accompanied by increased errors on *ω* and *κ*.

Thus, overall speedup on the simple M0 model can be explained by average observed codons count and by alignment length (Fig. 8). The relationship between speedup and *ω*, *κ*, tree length and codon frequencies is effectively explained by a reduced size of the state space of the continuous-time Markov chain. Aggregation is thus all the more effective when sequence data are biased or when analyses contain closely related species, which is probably the case for many real multiple sequence alignments.

**Figure 8:**
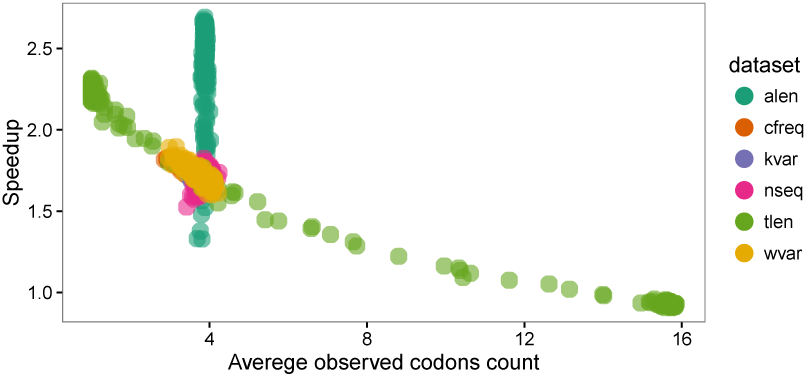
Speedup versus average codon count for M0 model. Each point represents one simulated alignment, dataset indicated by color. For the dataset with changing alignment length (`alen`), variation in speedup does not depend on the observed codon count (which does not vary significantly), but longer alignments lead to higher speedup, see Fig. 5.

### Branch-Site Model

Given the small speedup that we obtained for the aggregation on fixed positions, we implemented only the full aggregation mode for the branch-site model in FastCodeML. We then compared this new implementation with the standard FastCodeML. We see a slight increase in both false positives and true positives with aggregation (Tables 1, 2). Overall, ROC curves show that the performance of FastCodeML in aggregation mode is similar to the full likelihood mode (Fig. 9). Thus any errors in estimation under aggregation seem to have very little impact on the Likelihood ratio test (LRT) used to test for the presence of positive selection with the branch-site model.

**Table 1:** Statistical performance of FastCodeML in normal and aggregated modes on simulated data. Numbers in the cells correspond to the number of performed tests. A single branch was tested per tree.

| Mode | True positives | True negatives | False positives | False negatives |
| --- | --- | --- | --- | --- |
| normal | 551 | 973 | 27 | 449 |
| aggregated | 562 | 970 | 30 | 438 |

**Table 2:**
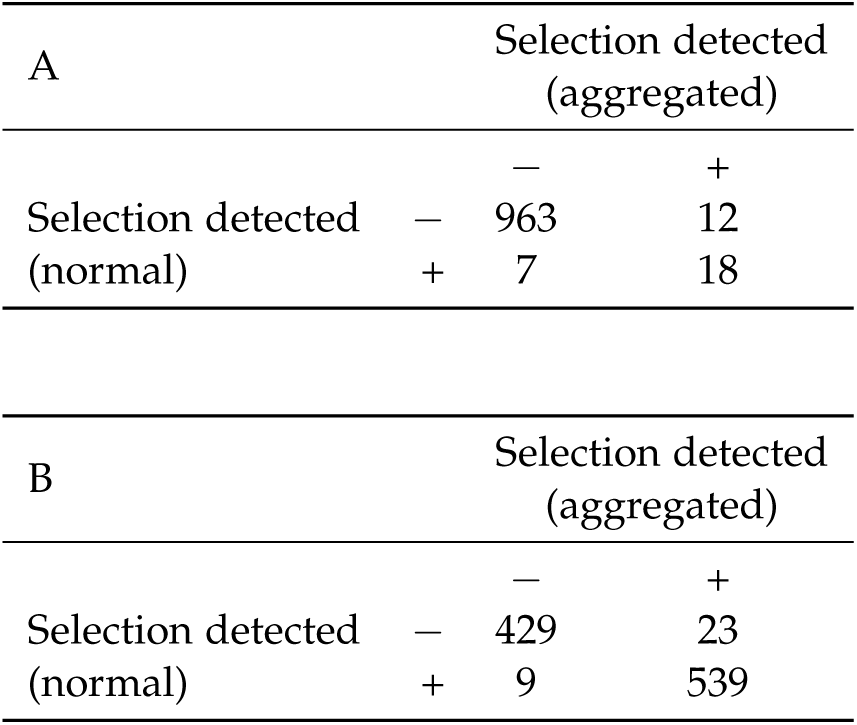
Statistical performance of FastCodeML on the simulated dataset. Numbers in the cells correspond to the number of performed tests. A single branch was tested per tree. A) Without positive selection; B) With positive selection.

**Figure 9:**
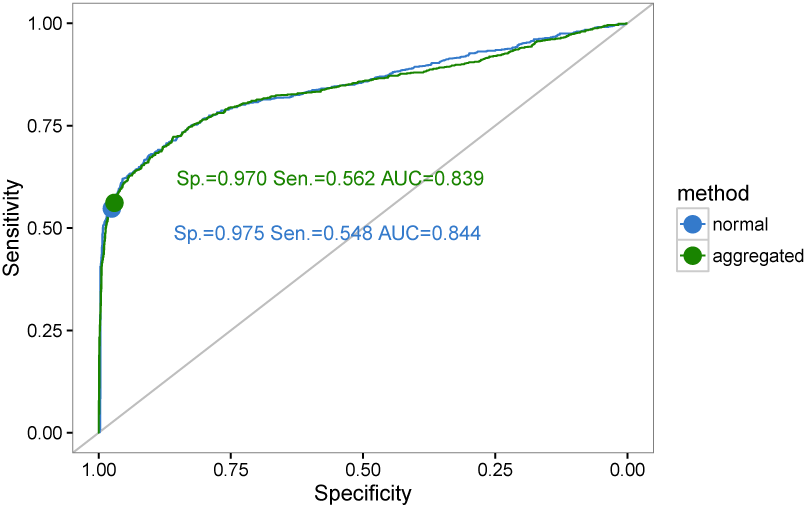
ROC (receiver operating characteristic) curves for FastCodeML in full likelihood and aggregated likelihood modes for the branch-site model simulations. Specificity, sensitivity and area under curve (AUC) indicated.

For *ω*_0_, *κ*, and *p*_1_, Pearson’s correlation coefficients between aggregated and full likelihood estimates are 0.9986, 0.9969 and 0.9735, respectively (Fig. 10). A lower correlation is observed for *p*_0_ and *ω*_2_ (0.9578 and 0.9109, respectively). Yet, these correlations are much higher than those obtained between the full likelihood estimate and simulated values: 0.35 for *p*_0_ and 0.20 for *ω*_2_.

**Figure 10:**
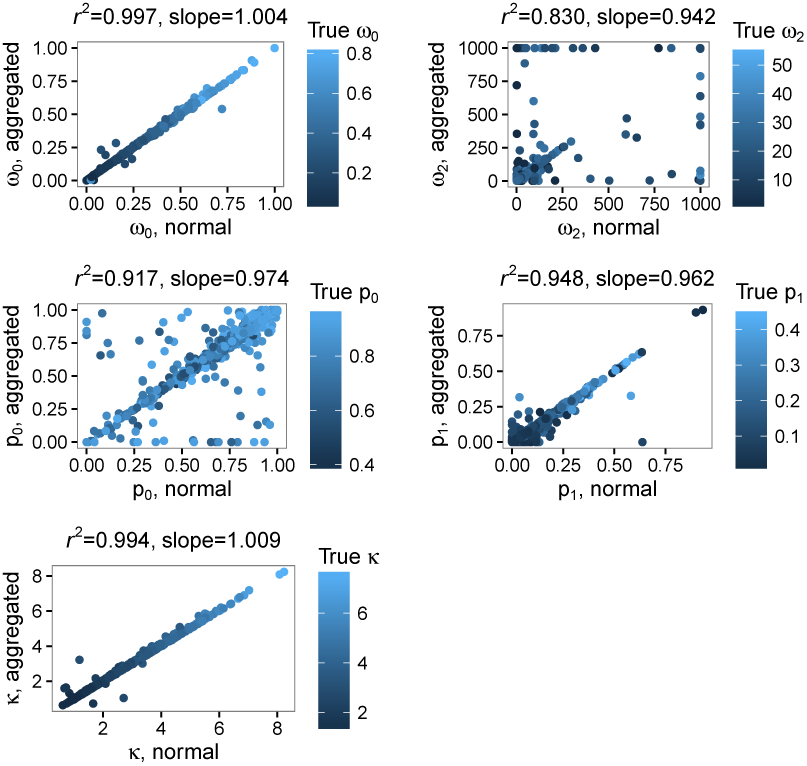
Correlation between aggregated and non aggregated parameter estimates for the branchsite model.

As with the M0 model, speedup is mostly affected by sequence length and tree length (Fig. 11) through their effects on observed codon counts (Fig. S19). We reached a maximum speedup of 4.4 fold per likelihood computation for the branch-site model.

**Figure 11:**
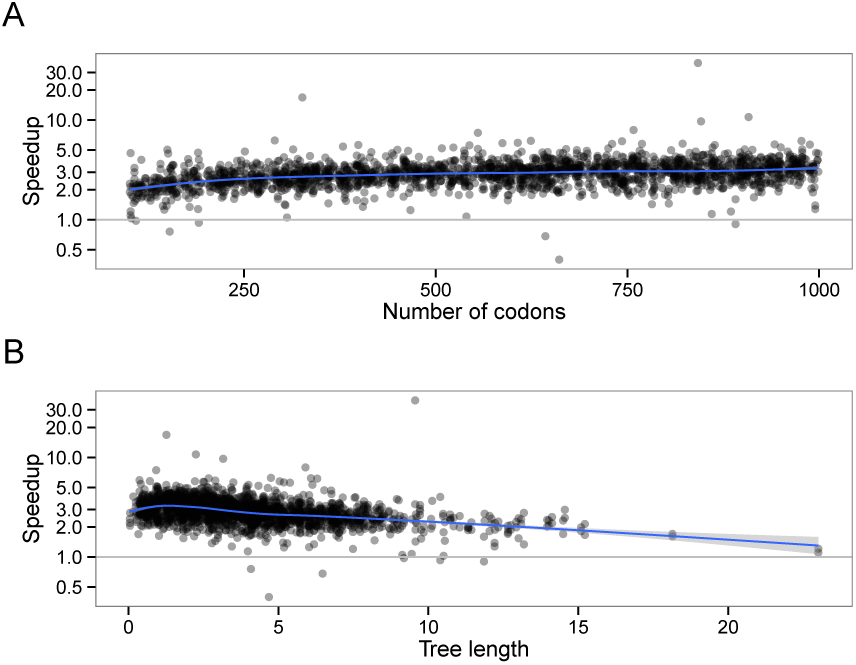
Effect of A) alignment length and B) tree length on the speedup, branch-site model.

The extended branch-site model violates several assumptions of the branch-site model (homogeneous synonymous rate, fixed *ω*_0_ and *ω*_2_ values). In those cases, where data is more complex than the model, the performance of the aggregated mode becomes slightly worse compared to the full likelihood mode S23, although it remains very close (AUC 0.812 *vs.* 0.818).

Finally, we used FastCodeML in normal and aggregated modes on a real dataset from Primates (Tables 3, S3). After correction for multiple testing (false discovery rate cutoff 0.05), 20 branches were identified to be under positive selection using full likelihood computations and 18 using aggregation, with 13 branches in common. We did not encounter multiple branches detected for an individual tree. The predictions are consistent between the two methods in 99.97% of the cases, which is higher than the consistency of 97.45% for the simulated data (Table 2). Aggregation gives a median speedup of 2.7 on this real dataset, confirming that real data can be sufficiently biased to make aggregation quite efficient.

**Table 3:**
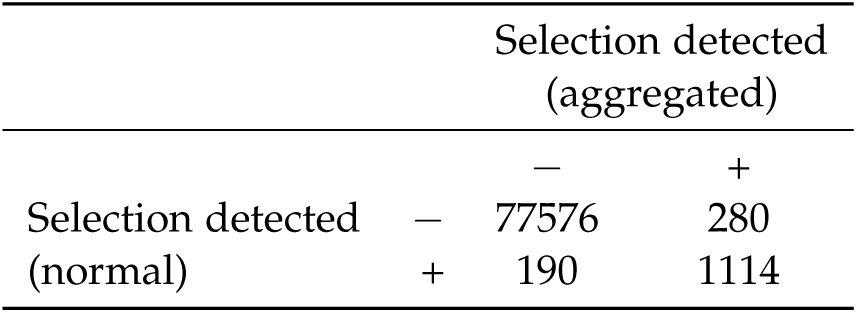
Statistical performance of FastCodeML on the Primates dataset. Detected selection in normal and aggregated modes of FastCodeML. Numbers in the cells correspond to the number of performed tests. Every non-terminal branch was tested.

## Discussion

We propose state aggregation as a technique for speeding up the computation of likelihood in a phylogenetic context. By reducing the size of the state space of the Markov process, aggregation accelerates the phase of tree pruning during the likelihood computation and, in some cases, the eigendecomposition of the transition rate matrix. We show that aggregation can be applied to the likelihood calculation of two of the most commonly used codon models. It can also be used for other types of models (see below), in both maximum likelihood and Bayesian frameworks.

The speedup for codon models depends on the alignment length and the observed codon counts, the latter being mostly affected by the tree length (Fig. 8, S19).

These effects are especially strong with the M0 model, because the likelihood optimizer uses a fixed number of iterations. A similar trend is observed with a variable number of iterations, but with increased stochasticity (Fig. S20). In general, state aggregation does not appear to have a systematic influence on the total number of iterations (Fig. S21). The total run-time is therefore proportional to the likelihood computation time.

Alignment length and observed codon counts have a similar effect on a speedup for the branch-site model. We see more explicitly the dependency if we normalize for the number of likelihood function computations (Fig. S22).

The most time consuming stages of the likelihood computation are matrix exponentiation and tree pruning. FastCodeML uses highly optimized algorithms to do matrix exponentiation (Schabauer *et al.,* 2012) and state aggregation improves the time to perform the tree pruning steps of the likelihood calculations (Fig. S1 A, B).

While the dependency of the speedup on the alignment length and the codon counts make intuitive sense, we can understand it in more details by considering the steps of the likelihood computation. Let us consider the computation time of the total likelihood (Fig. S1 A):

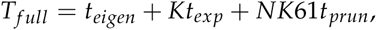

where *t*_*eigen*_ is the time to decompose the instantaneous rate matrix, *t*_*exp*_ is the time to exponentiate the rate matrix for each internal node, *t*_*prun*_ is the time to compute the partial likelihood vector per internal node per position, *K* is the number of internal nodes and *N* is the number of positions in the alignment. The number of states is 61 for Markov chains modeling codon sequences.

Similarly for the state aggregation (Fig. S1 B):

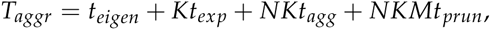

where *t*_*agg*_ is the matrix aggregation time per internal node per position, and 61 is replaced by M, the number of states after aggregation. For a given branch and site combination, the aggregation time is comparable to the time spent computing a single element of the partial likelihood vector. In the full mode, 61 elements of the vector should be computed. The gain of computing time observed with the aggregation methods comes from the need to do a single aggregation step, which is fast, followed by the computation of *M* (< 61) vector elements.

Aggregation speedup is thus:

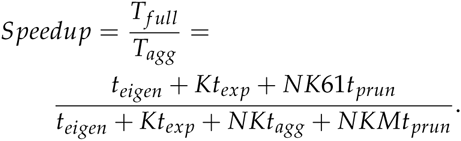

Generally performance is limited by eigen-decomposition and pruning, so we can approximate speedup as:

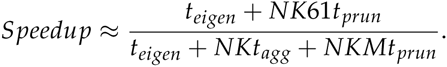

This representation gives a clear explanation for the dependency of the speedup on the alignment length and the observed codon counts. Increasing the alignment length causes a weaker effect on the non-accelerated eigendecomposition phase, which results in a more efficient acceleration. In contrast, a higher codon diversity in each alignment position increases the number of states in the aggregated Markov process (*M*), thus reducing the advantage of the aggregated process relative to the full one.

Not only does aggregation provide diminishing speedup with longer trees (more observed states, larger *M*), it also introduces a bias in the parameter estimation for extremely long trees. Consequently, for trees longer than 100 expected substitutions per position it is not practical to use state aggregation: biased results would be obtained without any significant speedup. In practice, however, extremely long trees are rare, for example in the Selectome database 99% of the trees has total length below 18 expected substitutions per position.

State aggregation can be applied either to the probability matrix *P* or to the instantaneous matrix *Q* (Fig. S1 B, C). In this work we were focused on applying aggregation to the probability matrix *P*. In this case (Fig. S1 B), aggregation is applied after exponentiation and must be performed for every position independently. The performance improvement is therefore achieved during the tree pruning phase. In the case of the matrix *Q* (Fig. S1 C), aggregation is applied prior to the exponentiation. This leads to smaller dimensions of *P* matrices, but eigendecomposition and exponentiation have to be performed for every position independently, since those positions will differ in the states aggregated. Moreover, aggregation of the matrix *Q* is expected to introduce more bias that will accumulate along the branches. Aggregation performed on the *Q* matrix will discard differences in substitution trajectories passing through unobserved states. There will thus be an accumulation of the error during both exponentiation and pruning phases. Aggregation done after the exponentiation phase only introduces error during the tree pruning phase. Preliminary results do not show an advantage of aggregating the matrix *Q* for codon models (not shown). A solution might be to perform a “softer” aggregation on clusters of sites with similar patterns of codons. This would be done by first clustering alignment positions and then producing aggregated instantaneous rate matrices for each cluster. This should diminish the bias and allow to exponentiate a smaller number of *Q* matrices than for the aggregation per site, while still computing on smaller *Q* matrices than in non aggregated mode. It is also possible that aggregation of the *Q* matrix could be more useful for other types of models, especially those with large instantaneous rate matrices, such as co-evolution models (Dib *et al.,* 2014). Finally, a second round of aggregation might be performed after the exponentiation in order to speedup the tree pruning stage (Fig. S1 D). The computational and statistical performance of such approaches has yet to be investigated.

It is also possible to implement aggregation on a subset of the data only. In our case, we chose an extreme situation and aggregated only the most conserved positions. The result was a large loss in speedup relative to aggregation on all positions without any gain in accuracy. But there might be other cases where aggregation on a subset of data only makes most sense in terms of the cost (accuracy) — benefit (speedup) trade-off. Moreover, there are multiple ways to perform the aggregation itself. Here, we collapsed all of the codons or amino-acids which are not observed at the position of the alignment. It is also possible to use other approaches to aggregation, e.g. aggregate all the codons reachable by more than a single mutation (or follow the amino-acids similarity properties). Models of amino acid substitutions have been derived from codon-based Markov models by aggregating codons separated by only synonymous substitutions. These models were, however, not built nor evaluated for computational efficiency (Yang *et al.,* 1998; Ren *et al.,* 2005; Susko and Roger, 2007; Kosiol and Goldman, 2011). Less aggressive aggregation shows increase in the accuracy at the price of reduced speedup, although, in our tests, accuracy was already good with the aggressive aggregation.

The combined use of both aggregated and non-aggregated modes in the same analysis could be efficient in several scenarios. First, aggregation could be used during likelihood maximization, but the final likelihood value computed without aggregation, providing a more accurate value. Second, aggregation could be used to obtain a starting point for non-aggregated likelihood maximization. Third, aggregation could be used in a preprocessing step to detect datasets of interest (e.g., gene families with a signal of positive selection). These datasets could then be analyze with full likelihood to get an accurate estimation of the parameters and model comparison. Finally, aggregation could be used during the burn-in period in a Bayesian approach (e.g. MCMC). There are probably other scenarios where aggregation can provide a faster estimation of likelihood within a more complex analysis.

For the specific case of the branch-site model, we have tested the second scenario of using aggregation as a starting point and we do not obtain a significant speedup (Fig. S24).

Aggregation can also have an impact on memory usage. Aggregation on the probability matrix *P* will reduce the size of the partial likelihood vectors (by a factor of *r* = 61/*N*_*states*_). Additionally the *Q* matrix aggregation reduces the size of the *P*-matrices (by a factor of *r*^2^). On the other hand, the actual improvement strongly depends on the details of the implementation, as partial likelihood vectors and probability matrices can be reused in a number of ways. Our implementation did not focus on the reduction of memory footprint and we thus do not discuss this aspect further.

Obviously, state aggregation in phylogeny and evolution is not limited to the branch-site and M0 codon models. First, it is universally applicable to Markov process-based codon models, such as the commonly used M1a/M2a, M8a/M8 (Wong *et al.,* 2004), aBSREL (Smith *et al.,* 2015), RELAX (Wertheim *et al.,* 2014), or any other GY94 (Goldman and Yang, 1994) or MG94-based (Muse and Gaut, 1994) model. Second, it is not limited to codon models. Given a trade-off between per-position matrix aggregation slowdown and tree pruning speedup, aggregation is unlikely to give a significant performance improvement for models with a small number of states (e.g., nucleotide models). But even for amino acids models we can expect some degree of speedup. In contrast, we expect state aggregation to provide a significant performance improvement for the models with a large number of states, such as amino acid coevolution models that can include up to 400 states (Yeang and Haussler, 2007; Dib *et al.,* 2014).

The aggregation of states in a Markov process is a powerful technique used in a variety of fields including computational biology, such as protein network interaction analysis (Petrov *et al.,* 2012), reaction modeling (Ullah *et al.,* 2012), single molecule photobleaching (Messina *et al.,* 2006), or disease-progression models (Regnier and Shechter, 2013). Its application to phylogenetic models has not been systematically studied, although it has been implemented in some software (Lartillot and Philippe, 2004, e.g. PhyloBayes;). This is, to our knowledge, the first systematic study of state aggregation biases and computational efficiency for molecular evolution.

In conclusion, we demonstrate that state aggregation is a powerful method which improves computational performance of codon-based models, with little cost in accuracy. State aggregation is not limited to codon models, and we expect it to be useful for a large variety of phylogenetic models and methods.

## ACKNOWLEDGMENTS

We would like to thank Bastien Boussau, Christophe Dessimoz, Stephane Guindon and anonymous referees for useful comments. The computations were performed at the Vital-IT (http://www.vital-it.ch) centre for high-performance computing of the Swiss Institute of Bioinformatics.

## References

Aldous, D. and Fill, J.A. (2002). Reversible markov chains and random walks on graphs. Unfinished monograph.

Cunningham, F. et al (2015). Ensembl 2015. Nucleic acids research, 43(D1), D662–D669.

de Koning, A.J. et al (2012). Phylogenetics, likelihood, evolution and complexity. Bioinformatics, 28(22), 2989–2990.

Dib, L. et al (2014). Evolutionary footprint of coevolving positions in genes. Bioinformatics, 30(9), 1241–1249.

Felsenstein, J. (1973). Maximum likelihood and minimum-steps methods for estimating evolutionary trees from data on discrete characters. Systematic Zoology, 22(3), 240–249.

Felsenstein, J. (1981). Evolutionary trees from DNA sequences: a maximum likelihood approach. Journal of molecular evolution, 17(6), 368–376.

Gladstein, D.S. (1997). Efficient incremental character optimization. Cladistics, 13(1), 21–26.

Goldman, N. and Yang, Z. (1994). A codon-based model of nucleotide substitution for protein-coding DNA sequences. Molecular biology and evolution, 11(5), 725–736.

Goloboff, P.A. (1993). Character optimization and calculation of tree lengths. Cladistics, 9(4), 433–436.

Guindon, S. and Gascuel, O. (2003). A simple, fast, and accurate algorithm to estimate large phylogenies by maximum likelihood. Systematic biology, 52(5), 696–704.

Guindon, S. et al (2010). New algorithms and methods to estimate maximum likelihood phylogenies: assessing the performance of PhyML 3.0. Systematic biology, 59(3), 307–321.

Hillston, J. (1995). Compositional markovian modelling using a process algebra. In Computations with Markov chains, pages 177–196. Springer.

Hordijk, W. and Gascuel, O. (2005). Improving the efficiency of SPR moves in phylogenetic tree search methods based on maximum likelihood. Bioinformatics, 21(24), 4338–4347.

Huelsenbeck, J.P. et al (2001). MRBAYES: Bayesian inference of phylogenetic trees. Bioinformatics, 17(8), 754–755.

Kemeny, J. and Snell, J. (1983). *Finite Markov Chains: With a New Appendix “Generalization of a Fundamental Matrix”*. Undergraduate Texts in Mathematics. Springer New York.

Kosiol, C. and Goldman, N. (2011). Markovian and non-markovian protein sequence evolution: aggregated markov process models. Journal of molecular biology, 411(4), 910–923.

Lartillot, N. (2006). Conjugate gibbs sampling for bayesian phylogenetic models. Journal of computational biology, 13(10), 1701–1722.

Lartillot, N. and Philippe, H. (2004). A bayesian mixture model for across-site heterogeneities in the amino-acid replacement process. Molecular biology and evolution, 21(6), 1095–1109.

Messina, T.C. et al (2006). Hidden markov model analysis of multi-chromophore photobleaching. The Journal of Physical Chemistry B, 110(33), 16366–16376.

Moretti, S. et al (2014). Selectome update: quality control and computational improvements to a database of positive selection. Nucleic acids research, 42(D1), D917–D921.

Murrell, B. et al (2012). Detecting individual sites subject to episodic diversifying selection. PLoS Genet, 8(7), e1002764–e1002764.

Muse, S.V. and Gaut, B.S. (1994). A likelihood approach for comparing synonymous and nonsynonymous nucleotide substitution rates, with application to the chloroplast genome. Molecular Biology and Evolution, 11(5), 715–724.

Nguyen, L.T. et al (2014). IQ-TREE: A fast and effective stochastic algorithm for estimating maximum-likelihood phylogenies. Molecular Biology and Evolution, page msu300.

Petrov, T. et al (2012). Model decomposition and stochastic fragments. Electronic Notes in Theoretical Computer Science, 284, 105–124.

Phillips, M.J. et al (2004). Genome-scale phylogeny and the detection of systematic biases. Molecular biology and evolution, 21(7), 1455–1458.

Proux, E. et al (2009). Selectome: a database of positive selection. Nucleic acids research, 37(suppl 1), D404–D407.

Regnier, E.D. and Shechter, S.M. (2013). State-space size considerations for disease-progression models. Statistics in medicine, 32(22), 3862–3880.

Ren, F. et al (2005). An empirical examination of the utility of codon-substitution models in phylogeny reconstruction. Systematic Biology, 54(5), 808–818.

Rodrigue, N. et al (2008). Uniformization for sampling realizations of markov processes: applications to bayesian implementations of codon substitution models. Bioinformatics, 24(1), 56–62.

Ronquist, F. (1998). Fast Fitch-parsimony algorithms for large data sets. Cladistics, 14(4), 387–400.

Rubinstein, N.D. et al (2011). Evolutionary models accounting for layers of selection in protein-coding genes and their impact on the inference of positive selection. Molecular biology and evolution, 28(12), 3297–3308.

Schabauer, H. et al (2012). SlimCodeML: an optimized version of CodeML for the branch-site model. In 2012 IEEE 26th International Parallel and Distributed Processing Symposium Workshops & PhD Forum, pages 706–714. IEEE.

Smith, M.D. et al (2015). Less is more: An adaptive branch-site random effects model for efficient detection of episodic diversifying selection. Molecular biology and evolution, 32(5), 1342–1353.

Stamatakis, A. (2014). RAxML version 8: a tool for phylogenetic analysis and post-analysis of large phylogenies. Bioinformatics, 30(9), 1312–1313.

Stamatakis, A. et al (2005). RAxML-III: a fast program for maximum likelihood-based inference of large phylogenetic trees. Bioinformatics, 21(4), 456–463.

Stamatakis, A.P. et al (2004). New fast and accurate heuristics for inference of large phylogenetic trees. In Parallel and Distributed Processing Symposium, 2004. Proceedings. 18th International, page 193. IEEE.

Storey, J.D. et al (2004). Strong control, conservative point estimation and simultaneous conservative consistency of false discovery rates: a unified approach. Journal of the Royal Statistical Society: Series B (Statistical Methodology), 66(1), 187–205.

Susko, E. and Roger, A.J. (2007). On reduced amino acid alphabets for phylogenetic inference. Molecular biology and evolution, 24(9), 2139–2150.

Swofford, D.L. and Olsen, G.J. (1990). Phylogeny reconstruction. In D. Hillis and C. Moritz, editors, Molecular Systematics, pages 411–501. Sinauer Associates, Sunderlands, Massachusetts.

Ullah, G. et al (2012). Simplification of reversible markov chains by removal of states with low equilibrium occupancy. Journal of theoretical biology, 311, 117–129.

Valle, M. et al (2014). Optimization strategies for fast detection of positive selection on phylogenetic trees. Bioinformatics, page btt760.

Vera-Ruiz, V.A. et al (2014). Statistical tests to identify appropriate types of nucleotide sequence recoding in molecular phylogenetics. BMC bioinformatics, 15(Suppl 2), S8.

Vilella, A.J. et al (2009). EnsemblCompara GeneTrees: Complete, duplication-aware phylogenetic trees in vertebrates. Genome research, 19(2), 327–335.

Wertheim, J.O. et al (2014). Relax: detecting relaxed selection in a phylogenetic framework. Molecular biology and evolution, page msu400.

Wong, W.S. et al (2004). Accuracy and power of statistical methods for detecting adaptive evolution in protein coding sequences and for identifying positively selected sites. Genetics, 168(2), 1041–1051.

Yang, Z. (2007). PAML 4: phylogenetic analysis by maximum likelihood. Molecular biology and evolution, 24(8), 1586–1591.

Yang, Z. et al (1998). Models of amino acid substitution and applications to mitochondrial protein evolution. Molecular Biology and Evolution, 15(12), 1600–1611.

Yeang, C.H. and Haussler, D. (2007). Detecting coevolution in and among protein domains. PLoS Comput Biol, 3(11), e211.

Zhang, J. et al (2005). Evaluation of an improved branch-site likelihood method for detecting positive selection at the molecular level. Molecular biology and evolution, 22(12), 2472–2479.

